# CRISPR-DA: Adapting CRISPR gRNA design for detection assays

**DOI:** 10.64898/2025.12.05.692701

**Authors:** Carl Schmitz, Jacob Bradford, Dimitri Perrin

## Abstract

CRISPR-Cas systems offer a viable alternative to traditional detection and diagnosis methods. However, their effectiveness relies heavily on the selection of appropriate guide RNA sequences. Existing gRNA design tools were primarily developed for gene editing and are not always directly applicable to CRISPR-based detection assays. In particular, alignment-based methods are still used to estimate gRNA specificity, even though they can miss a substantial portion of off-target sites. In this work, we introduce CRISPR-DA, a CRISPR gRNA design tool for detection assays. We show that it provides a better assessment of gRNA specificity than BLAST, which detected only 33.27 % and 0.43 % of cross-species off-targets in two datasets. Additionally, CRISPR-DA ran two and six times faster than BLAST on these datasets, respectively. Our method incorporates advances from gene-editing guide RNA design tools, including uncertainty-informed guide RNA design, to improve the selection of guides with high on-target activity. CRISPR-DA is available at https://github.com/bmds-lab/CRISPR-DA.

## Introduction

Accurate species detection is critical for several fields. For instance, viruses and infectious diseases pose a significant threat to public health, as highlighted by recent outbreaks of SARS-CoV-2 and, historically, by the Spanish flu, swine flu, bird flu, Ebola, as well as the Zika virus (1). Their timely detection is crucial to limiting spread and enabling the application of appropriate treatments (2).

Similar risks exist in food security, where viruses have the potential to infect or devastate key crops such as wheat, rice, maize, potato, and soybeans (3). Not only does this represent a significant economic concern, but it also endangers global food security (4). Detection and diagnosis techniques serve as a powerful tool to help manage the threat of plant viruses (5, 6).

Detection methods can also be applied to identify genomic material in environmental DNA (eDNA) (7). eDNA can be used to monitor endangered and rare species (8) or detect invasive ones (9).

In all these contexts, detection is the vital first step: we cannot effectively control human or crop pathogens without the ability to detect them. Likewise, we cannot conserve endangered species or manage invasive ones without knowing their spatial distributions.

Currently, polymerase chain reaction (PCR) methods remain the gold standard for detecting genomic material in samples of interest due to their high precision and minimal error rates (10, 11). Metagenomic Next-Generation Sequencing (mNGS) and Nanopore Third-Generation Sequencing (NTS) have been highlighted for their high sensitivity and rapid sampling process, but they lack validation guidelines and are more expensive than other methods (12). CRISPR-Cas systems offer a promising, low-cost, and easily deployable alternative for detecting specific genomic material (13).

Although originally discovered as an adaptive immune response in many bacteria and archaea (14), CRISPR-Cas systems have since been used as gene-editing tools (15). The key component of the CRISPR-Cas system is the CRISPR-associated (Cas) protein, which can target specific DNA sequences that match a programmable guide RNA (gRNA).

The DNA recognition property of CRISPR-Cas systems has been harnessed to guide other biological functions to specific DNA locations (16–18). One such application is the development of novel detection methods (19). CRISPR-Cas systems used for diagnostic purposes typically focus on Cas proteins such as Cas9, Cas12, and Cas13 (20). They have been used successfully to detect a range of viruses and diseases, including the Zika virus (21), Mycobacterium tuberculosis (22), SARS-CoV-2 (23), and several plant viruses such as Potato Virus X (24). In addition to the detection of viruses and diseases, CRISPR-Cas systems have also been applied to species identification (25) and species detection using eDNA (26).

Some commonly used CRISPR-Cas-based detection methods include DETECTR (27), SHERLOCK (28), and SHERLOCKv2 (29). These have the advantage of being faster, cheaper, and simpler than PCR, NTS, and mNGS methods (30). These advantages make CRISPR-Cas-based detection well-suited for field testing.

Regardless of the Cas protein or application, one common, non-trivial challenge is the selection of an appropriate gRNA to target the sequence of interest. In a gene-editing context, this selection is driven by two properties. The first is to consider the on-target efficiency, defined as the frequency at which the intended genomic modification is successfully achieved. The second property relates to specificity and the risk of inducing off-target modifications at sites that have a sequence similar enough (containing one to four mismatches) to be erroneously targeted. Both properties remain relevant in a detection context, where they directly relate to the sensitivity and specificity of the assay.

The specificity of the selected gRNA is important, as potential off-target activity is recognised as a major limitation of CRISPR-Cas-based detection and diagnosis (30, 31). Off-target activity would lead to false positives, increasing the error rate. Additionally, the on-target efficiency is important to ensure that the target site is consistently detected, thereby increasing sensitivity.

To compete with the gold standard PCR methods, CRISPRCas-based detection must ensure high precision with minimal errors through the careful selection of gRNAs that maximise on-target efficiency while minimising off-target activity.

Guide selection has been extensively studied in a gene-editing context, and current practices bear little resemblance to initial approaches. For instance, early approaches to gRNA design relied on sequence alignment tools to detect off-target sites. However, sequence alignment tools fail to identify all off-target sites (32). Consequently, newer gRNA design tools for gene editing have adopted dedicated methods to achieve comprehensive off-target detection. The same challenge remains in the context of detection assays, and guide RNAs must exhibit high specificity. However, despite the advances made in a gene-editing context, gRNA selection for a detection or diagnosis context still often relies on BLAST (see (33)). This highlights the need for gRNA design tools for detection and diagnosis applications that can leverage developments made in a gene-editing context.

At a high level, the principles of on-target efficiency and off-target specificity from gene editing still apply to detection assays. They directly translate to the sensitivity and specificity of the assays. However, modifications are necessary to account for the distinct nature of this application. For example, in gene-editing applications, an off-target effect is a modification somewhere else *in the same genome*, whereas, for a detection assay, we need to avoid cross-reaction with genomic material from other strains/species. This requires an off-target assessment *across multiple genomes*. While the principles and approaches used in gRNA design tools developed for gene-editing are relevant, they need to be extended to the multi-genome context, and the lack of dedicated tools tailored for detection assays remains a key limitation.

Here, we present CRISPR-DA (CRISPR guide RNA design for detection assays), a comprehensive gRNA design pipeline developed for CRISPR-Cas9-based detection and diagnostic applications. Our method incorporates an enhanced off-target scoring approach, demonstrated to be the fastest available, and ensures the accurate assessment of off-target risk. Additionally, the pipeline utilises a deep-learning ensemble that leverages uncertainty quantification to enable uncertainty-informed selection of gRNAs.

## Methods

### A. Off-target specificity

Off-target specificity has been identified as an area for improvement in CRISPR-Cas-based detection and diagnosis, regardless of the Cas protein used (30, 31). Here, we build upon the extensive work done on existing off-target analysis methods for Cas9.

Existing off-target analysis methods for gene-editing fall into two categories: alignment-based and score-based strategies (34). Sequence alignment tools such as BLAST, BLAT, Bowtie, and BWA are often used to identify potential offtarget sites. However, different off-target sites do not carry the same level of risk. For example, off-target sites identified for one gRNA may not present the same risk as those for another. Alternatively, score-based strategies, such as the MIT (35) and the Cutting Frequency Determination (CFD) (36) methods, quantify off-target risk by analysing mismatch position and, in the case of CFD, mismatch type.

However, commonly used sequence alignment tools, such as BLAST, have been shown to miss a significant number of offtarget sites, particularly those containing three to four mismatches (32). This limitation arises from the alignment algorithms themselves rather than the Cas protein used. As a result, gRNA design tools for gene-editing have developed dedicated off-target analysis methods, generally focused on Cas9.

In the context of detection and diagnosis, incomplete off-target detection can lead to unintended activity. This could lead to false positives, increasing the error rate. This is an issue that can be avoided with accurate off-target identification.

A key distinction is that gene-editing studies typically assess off-target sites within a single genome, whereas detection and diagnostic assays require evaluation across multiple genomes. Therefore, in order to utilise existing off-target analysis methods designed for gene-editing, they must be run across multiple genomes to assess each gRNA’s specificity relative to its target genome.

Furthermore, off-target analysis is computationally intensive, influenced primarily by genome size and the number of gRNAs analysed. As highlighted by (37), many gRNA design tools are limited by extended run times. This issue is likely to persist and even worsen when analysing multiple genomes as required for gRNA design in detection and diagnosis.

Therefore, when selecting the Cas9 off-target analysis method for our tool, it was essential that the method operate efficiently to enable off-target analysis across multiple genomes. Additionally, it needed to accurately detect all off-target sites to minimise the risk of false positives.

Our previous gRNA design tool for gene-editing using Cas9, Crackling++ (38), utilised a modified version of Inverted Signature Slice Lists (ISSL) (39) to rapidly identify all off-target sites and calculate MIT and CFD scores. The full method is detailed in (38); however, the key points are summarised below. The first step involves identifying all off-target sites in the genome of interest. Next, the off-target sites were arranged into the ISSL data structure based on partial matches, such that all off-target sites in each list share the same partial match. To identify off-target sites for a given gRNA, only the lists sharing partial matches with that gRNA are searched. This resulted in two key outcomes: first, it greatly reduced the search space by excluding all off-target sites that have no similarity to the gRNA. Second, it guaranteed that all off-target sites with one to four mismatches exist in at least one list due to the configuration of the partial matches. Crackling++ is among the fastest and most scalable methods available. Accordingly, the Crackling++ off-target scoring module was integrated into CRISPR-DA.

### B. Benchmark

A benchmark was conducted to evaluate the performance of the Crackling++ off-target scoring module relative to BLAST in detecting Cas9 off-targets. BLAST was selected because Li et al. (32) reported that it outperformed other sequence alignment tools, although it still failed to identify all off-target sites. Another reason for selecting BLAST was its continued use in gRNA design for detection and diagnosis, as demonstrated by studies such as (33). The benchmark included two different datasets, which were created to simulate two distinct use cases.

The first dataset focuses on detecting one specific virus among other viral genomes. To create this dataset, a list of complete viral genomes was downloaded from the NCBI. To ensure computational feasibility, the first 2,001 viral genomes (one target and 2,000 potential off-target viruses) were selected. To ensure enough data were generated for later comparison, the viral genome with the highest number of gRNAs was designated as the target. From the total of 57,381 gRNAs identified, the first five batches of 10,000 gRNAs were sequentially selected, resulting in 50,000 gRNAs. The full list of viral genomes can be found in Supplementary Table 1.

**Table 1.** The number of gRNA above the acceptable threshold of 70 defined by Crackling++. The rows are divided by off-target scoring methods and the columns by off-target detection methods, where C indicates the Crackling++ off-target scoring module, and B indicates BLAST

|  | Domestic Cattle |  | Emu |  | Great White Shark |  | Zebrafish |  | Southern Bluefin Tuna |  | Virus dataset (average) |  |
| --- | --- | --- | --- | --- | --- | --- | --- | --- | --- | --- | --- | --- |
|  | C | B | C | B | C | B | C | B | C | B | C | B |
| MIT | 1,802 | 9,718 | 3,691 | 9,815 | 1,017 | 9,767 | 4,161 | 6,667 | 5,571 | 9,798 | 50,000 | 50,000 |
| CFD | 5,055 | 9,995 | 7,507 | 9,998 | 3,064 | 9,995 | 8,414 | 9,872 | 7,843 | 9,989 | 50,000 | 50,000 |

The second dataset focuses on species identification, where the goal could be either distinguishing one species among closely related taxa or detecting its presence in eDNA. This dataset contains a small number of large genomes with the goal of species identification. The yellowfin tuna (GenBank assembly GCA_914725855.2) was selected as the target genome, with five off-target species of varying genetic similarity included. These were the southern bluefin tuna (GenBank assembly GCA_910596095.1), zebrafish (GenBank assembly GCA_000002035.4), and great white shark (GenBank assembly GCA_017639515.1), along with more distantly related species such as domestic cattle (GenBank assembly GCA_002263795.4) and the emu (GenBank assembly GCA_036370855.1). From the target species, the first 10,000 gRNAs were extracted and used for the benchmark.

The benchmark also included guides reported in existing literature for CRISPR–Cas9 lateral flow assays against the virus dataset. This benchmark would use two guides reported by (40) to detect the envelope and ORF1ab genes of SARS-CoV-2 and two guides from (41) used to detect Listeria monocytogenes (Lm) and African swine fever virus (ASFV). These studies were selected because they reported high detection accuracy but provided limited discussion of potential off-target effects.

Next, all required files were built for BLAST (V2.9.0) and the Crackling++ off-target scoring module. All commands were run concurrently using Python’s multiprocessing module. The benchmark was conducted on a Linux-based machine running an Intel Core i7-5960X (3.0 GHz) processor, equipped with 32 GB of memory and configured with 32 GB of swap space.

The following command was used to run the Crackling++ off-target scoring module: `ISSLScoreOfftargets` {`issl-index.issl}` {`guides.fa`}`4 0 and` > {`output.txt`}.

The following command was used for BLAST: `blastn -task blastn-short -ungapped` `-num_threads 32 -db` <`ref`> `-query` <`guides.fa`> `-out` <`output.txt`> `-evalue` `1 -outfmt “6 qacc stitle sstrand qstart` `qend sseq sstart send”`. ‘blastn’ was selected as the comparison involves two nucleotide sequences. The ‘-task blastn-short’ option was used as it is optimised for sequences shorter than 50 bases, making it suitable for CRISPR-Cas9 gRNAs, which are 20 bases long. The ‘-ungapped’ option was included to avoid gapped alignments, as off-target analysis focuses on pairwise mismatches. The ‘-evalue’ was set to 1 to ensure all results were included. Finally, the ‘-outfmt’ options (sequence strand (sstrand), sequence start (sstart), sequence end (send), etc.) were specified to locate the alignment within the overall sequence and determine whether a valid off-target site was detected. Once the benchmark was completed, the MIT and CFD off-target scores were calculated using the off-target sites reported by BLAST. Because the Crackling++ off-target scoring module computes scores rather than directly reporting sites, a modified version was implemented to output both total and unique off-target counts along with their corresponding scores.

## Results

### C. Off-target count

Table 2 summarises the off-targets reported by BLAST and the Crackling++ off-target scoring module for both datasets. Because the Crackling++ off-target scoring module identifies all off-targets, its reported counts and corresponding MIT and CFD scores were treated as the ground truth.

**Table 2.** The number of off-targets detected for all datasets divided by detection method.

|  | ISSL |  | BLAST |  |
| --- | --- | --- | --- | --- |
|  | Total | Unique | Total | Unique |
| Virus | 2,306,066 | 2,174,767 | 724,301 | 693,488 |
| Animal | 23,947,048 | 15,077,330 | 103,971 | 45,005 |
| Assay | 282 | 272 | 101 | 101 |

BLAST detected only 31.41 % of off-targets in the virus dataset, 0.43 % in the animal dataset, and 35.82 % for the CRISPR-Cas9 assay guides. Out of the 100 million BLAST runs for the virus dataset (50,000 gRNAs against 2,000 genomes), 1.21 % (1,210,477) were incorrectly reported as lacking off-target sites. For the animal dataset, 50,000 runs (10,000 gRNAs against five genomes) showed that 58.46 % (29,230) were also incorrectly reported as lacking off-target sites. For the guides used in CRISPR-Cas9 lateral flow assays, a total of 8,000 runs (four guides against 2,000 genomes), 1.99 % (159) were incorrectly reported as having no off-target sites. This represents a substantial number of unidentified off-targets, leading to an underestimation of off-target risk. In turn, this could result in false positives for detection and diagnosis if off-target activity were to occur. To assess the effect of missing off-targets, we analysed the MIT and CFD scores. For the virus dataset, 1.29 % (1,291,708) of MIT scores and 1.16 % (1,160,897) of CFD scores were underestimated. Notably, three of the CFD scores and 307 of the MIT scores differed by more than 50 points, a substantial difference given that the scores range from 0 to 100. For the animal dataset, 100 % of MIT and CFD scores were underestimated, with 26.21 % (13,107) and 10.00 % (5,000) differing by more than 50 points, respectively. For the assay guides, 2.09 % (167) of MIT scores and 1.65 % (132) of CFD scores were different. However, for the assay dataset, all score differences were less than 5 points. Figure 1 shows a histogram of the MIT and CFD score differences for both datasets. The off-targets missed by BLAST resulted in widespread underestimation of MIT and CFD scores. Except for the CFD scores from the virus dataset, there were cases of extreme score differences (greater than 80 points). For the virus dataset, the majority of differences were minimal (less than 5 points), with very few scores differing by more than 5 points. For the assay dataset, all score differences were minimal (less than 5 points).

**Fig. 1.**
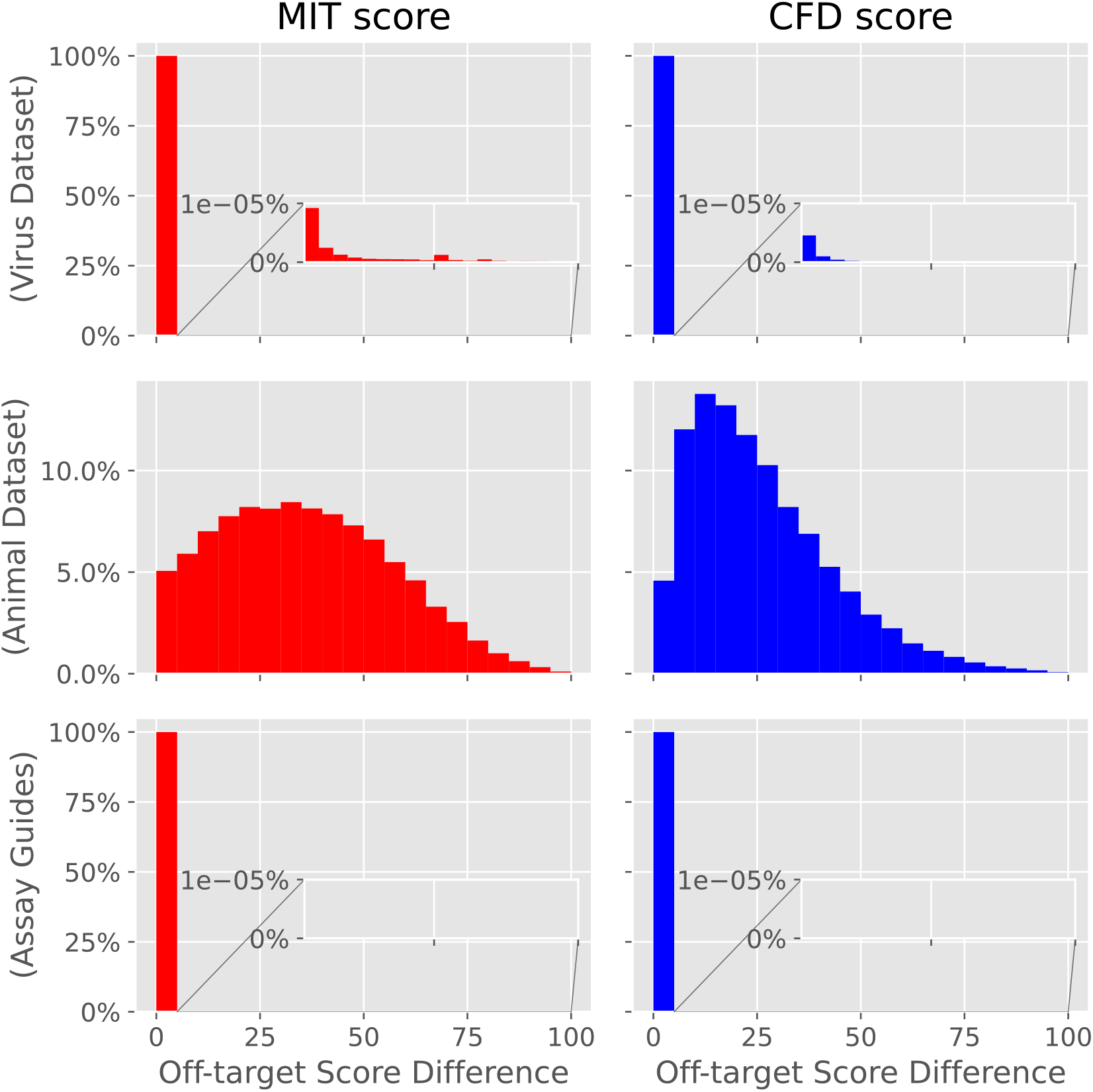
A histogram plot comparing the MIT and CFD score difference for both datasets when using BLAST instead of the Crackling++ off-target scoring module.

To evaluate the practical impact of missed off-targets on gRNA selection, a score threshold of 70 on the MIT and/or the CFD score, as utilised in Crackling++, was applied to determine which gRNAs would be accepted. Table 1 summarises the number of accepted gRNAs for both datasets. The virus dataset was averaged across all species, while the animal dataset was broken down by individual species.

For the virus dataset, both the Crackling++ off-target scoring module and BLAST accepted all gRNAs when averaged across all genomes. This highlights a limitation of using averages, as they can obscure outliers. To explore this further, the score threshold was applied to each individual genome in the virus dataset. BLAST would incorrectly calculate 514 MIT scores and 19 CFD scores to be above the threshold of 70. As only a small proportion of scores fell below the threshold, the overall quantitative impact on gRNA selection was minimal. For the animal dataset, there was a significant difference in the number of accepted gRNAs. Similar to the virus dataset, the MIT score was more affected by missed off-targets, resulting in a larger discrepancy than the CFD score. BLAST incorrectly calculated 29,523 MIT scores and 17,966 CFD scores as being above the threshold of 70. The southern bluefin tuna exhibited the least difference for both MIT and CFD scores. This was due to the high similarity between genomes, which resulted in a greater number of off-targets containing 0–2 mismatches, allowing them to be detected by BLAST.

For the assay guides, as all scores differed by less than 5 points, no difference was observed when using the threshold of 70.

### D. Mismatch count

As described by Li et al. (32), BLAST was able to identify highly similar off-targets (0–1 mismatches) but struggled as similarity decreased (2–4 mismatches). To verify this, the reported off-targets were binned by the number of mismatches they contained. This analysis was performed for both BLAST and the Crackling++ off-target scoring module results. The Crackling++ off-target scoring module uses the ISSL data structure, which, as summarised in Section A, employs partial matches to identify all off-target sites for a given gRNA. Because the Crackling++ off-target scoring module is guaranteed to report all off-targets, it was used as the 100 % baseline for comparison. In Table 3, we observed results similar to those reported in (32). BLAST was able to identify 97.01 % and 99.89 % of off-targets with 0 mismatches for the virus and animal datasets, respectively. However, detection rates declined sharply as the mismatch count increased, with fewer than 1 % of off-targets with three or four mismatches being detected in the animal dataset.

**Table 3.** The percentage of off-targets detected by BLAST compared to the Crackling++ off-target scoring module binned by mismatch count.

|  | 0 mismatch | 1 mismatch | 2 mismatch | 3 mismatch | 4 mismatch |
| --- | --- | --- | --- | --- | --- |
| Virus | 97.01 % | 88.55 % | 87.75 % | 64.41 % | 27.28 % |
| Animal | 99.89 % | 79.50 % | 9.52 % | 0.71 % | 0.04 % |
| Assay | 100.00 % | —* | 100.00 % | 23.08 % | 35.02 % |

\* Indicates that no off-targets with $x$ mismatches were present in that dataset.

### E. Run time

BLAST took an average of 403 seconds per batch of 10,000 gRNAs for the virus dataset and 4,089 seconds for the animal dataset. In contrast, the Crackling++ off-target scoring module was substantially faster, averaging 208 seconds per batch for the virus dataset and 544 seconds for the animal dataset. This represents approximately a twofold and sevenfold speed increase compared with BLAST for the virus and animal datasets, respectively.

## Discussion

### F. Missed off-targets

PCR methods remain the gold standard for detection and diagnosis due to their high accuracy and low error rates. While CRISPR-Cas-based detection and diagnosis offer the potential for a simpler, faster, more costeffective alternative, they require robust bioinformatics tools to achieve comparable accuracy and reliability.

As shown in Table 2, BLAST missed a substantial number of off-targets, particularly when analysing larger genomes. These undetected off-targets underestimate off-target risk and may result in false positives in practical applications. Furthermore, these results confirm that sequence alignment tools are inadequate for identifying all off-target sites. In contrast, gene-editing also requires that all off-target sites be detected, and as such, the Crackling++ off-target scoring module was designed to ensure comprehensive off-target identification. This highlights the advantages of purpose-built tools and how advances made for gRNA design for gene-editing can be adapted to create tools tailored for detection and diagnosis.

To investigate whether missed off-targets were isolated to specific species, we examined the number of gRNAs incorrectly reported as having no off-targets in the animal dataset. The results showed widespread under-reporting, with 8,133 gRNAs for zebrafish, 7,992 for the great white shark, 7,116 for the emu, and 5,864 for domestic cattle incorrectly identified as having no off-targets. The southern bluefin tuna was the only exception, with 125 gRNAs falsely reported as having no off-targets. This suggests that sequence alignment tools are capable of detecting highly similar off-target sites but still fail to capture all off-targets. These findings are consistent with the results in Table 3 and those reported by (32). To test the effect of missing a single off-target site, the maximum score drop was determined. This was achieved by generating all possible off-target combinations and calculating their corresponding scores. It was observed that the MIT score could decrease by approximately 50 points for one mismatch, 9.44 points for two, 2.54 points for three, and 1.37 points for four mismatches. For the CFD score, off-targets containing one to four mismatches could reduce the score by up to 0.99 points.

### G. Potential off-targets in CRISPR-Cas9 Assays

The benchmarking of guides used by (40) and (41) revealed, as shown in Table 2, off-target sites do exist and that relying solely on alignment-based methods such as BLAST would overlook a significant proportion of them. In contrast, Figure 1 demonstrates that the effect on the MIT and CFD scores was minimal. Notably, the MIT and CFD scores were developed to provide an improved metric for understanding off-target likelihood rather than for comprehensive detection.

To further investigate differences in off-target scores, the list of viruses incorrectly classified as free from off-targets was examined. In particular, the guide used to detect the envelope gene in SARS-CoV-2 would have missed off-targets in key viruses such as cowpox virus and monkeypox virus. The remaining misclassified viruses were primarily unrelated phages that are unlikely to be present. While the difference in scores was minimal, the potential effect of this variation in the context of detection and diagnosis remains unclear. Missed off-targets may increase assay error rates and result in false positives. It remains uncertain whether existing thresholds and scoring systems are directly applicable in this new context. Nevertheless, regardless of the specific scoring method or thresholds employed, accurate detection of all off-target sites remains paramount.

Furthermore, these results could be improved by selecting relevant viruses for off-target testing. The virus dataset used for benchmarking was constructed by randomly selecting viral genomes. Including potential high-risk genomes likely to be present in the tested samples would reduce irrelevant comparisons and increase the significance of any observed decrease in off-target scores. For example, when designing a guide to detect the foodborne pathogen Lm, the off-target genomes should include only foodborne pathogens.

### H. Runtime

gRNA design tools for gene-editing have developed specialised methods to improve off-target analysis and reduce runtimes for more efficient processing. However, due to the lack of gRNA design tools specifically built for detection and diagnosis, this research gap remains unaddressed. Efficient analysis is particularly important in this context, given the requirement to process data across multiple genomes.

Section E presents the runtimes for both datasets, showing that the Crackling++ off-target scoring module outperforms BLAST in both cases. This is significant because Crackling++ performs additional work by calculating both the MIT and CFD scores. BLAST only reports potential off-target sites and requires further processing to compute these scores. These results demonstrate the suitability and efficiency of the Crackling++ off-target scoring module for integration into CRISPR-DA. They also highlight how advances from geneediting gRNA design tools can be effectively applied to detection and diagnostic tools, offering reduced runtimes compared with general-purpose sequence alignment methods.

### I. Visualising results

CRISPR-DA calculates an average off-target score across all analysed genomes for each gRNA, using both the MIT and CFD metrics. However, relying on an average score may obscure outliers and misrepresent the true off-target risk. To address this limitation, we developed a phylogenetic-tree-based method for visualising off-target risk, enabling more informed decision-making in gRNA design.

This functionality was implemented using the web-based Interactive Tree of Life (iTOL) (42), allowing users to visualise potential off-target risks for each genome included in the analysis. To enable this, CRISPR-DA generates a phylogenetic tree of all analysed genomes, exported in Newick format, along with annotation files compatible with iTOL. On the iTOL website, users can interact with the phylogenetic tree. Users can also collapse any clade to view the average off-target score at different taxonomic levels (e.g. species or family), mouse over nodes for more information, and toggle between MIT and CFD scores. Finally, the results can be output to various image formats. See Figure 2 for an example of the image output.

**Fig. 2.**
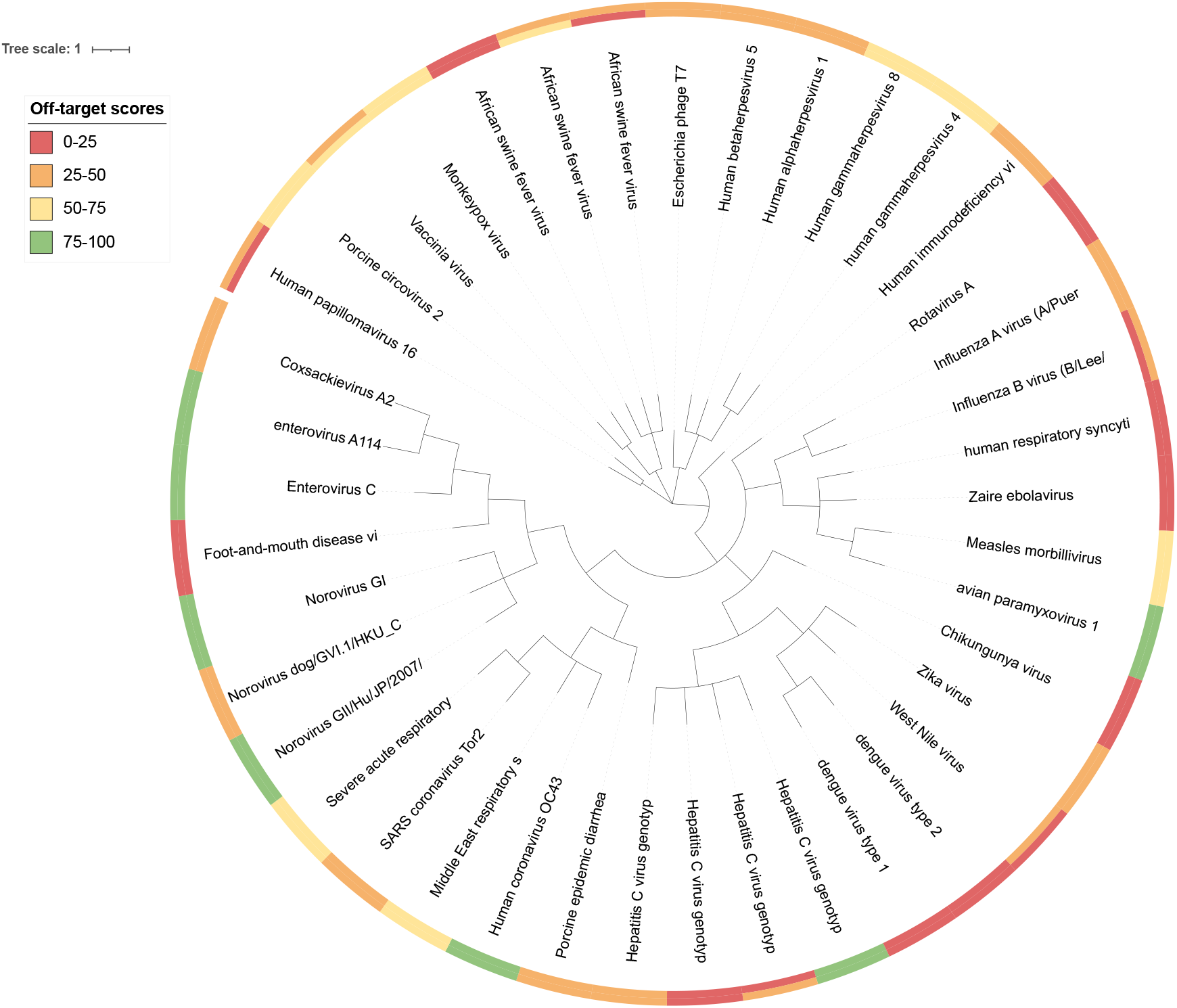
Interactive Tree of Life visualisation of CRISPR-DA results. A clade can be collapsed to summarise the scores of all descendants. Selecting a node reveals details like off-target counts. Ring segment colours indicate off-target risk severity; dual colours appear when MIT and CFD scores differ. The inner colour reflects the first imported score.

### J. On-target efficiency

In order to comprehensively design gRNAs, the on-target efficiency must also be assessed. When assessing gRNA on-target efficiency, there is little difference between the contexts of gene editing and detection or diagnosis. Candidate sites for both gene editing and detection exist within a single genome. Therefore, existing methods can be utilised for assessing gRNA on-target efficiency. Ensuring the use of gRNAs with high on-target efficiency increases the true positive rate and, in turn, the precision of CRISPR-Cas-based detection and diagnostic methods.

With the continued development of deep learning models such as CRISPRon (43), DeepSpCas9 (44), DeepHF (45), and DeepCRISPR (46), these tools have been reported to accurately predict gRNAs with high on-target efficiency. However, there is a risk in treating deep learning methods as black boxes without appropriate metrics to assess prediction quality (47, 48).

For CRISPR-DA, we aimed to leverage the enhanced performance of state-of-the-art deep learning models while avoiding black-box implementation. Accordingly, appropriate metrics were incorporated to critically evaluate model predictions. To meet these objectives, we selected our previously developed Cas9 on-target efficiency assessment method, CRISPRDeepEnsemble (49). CRISPRDeepEnsemble is a deep learning method that provides accurate predictions of gRNA on-target efficiency while also generating uncertainty metrics. These uncertainty metrics offer insight into prediction confidence, enabling uncertainty-informed gRNA selection.

CRISPRDeepEnsemble captures epistemic (model-based) uncertainty by employing an ensemble of models rather than a single model. Each model is trained with different initialisation parameters, producing slight prediction variations that reflect epistemic uncertainty. By aggregating predictions and samples from all ensemble members, we derive a robust prediction and a quantification of the overall prediction uncertainty.

To capture aleatoric (data-based) uncertainty, each CRISPRDeepEnsemble member predicts the parameters of a Beta distribution, rather than directly estimating the score. To compute the predicted score, we collect *N* samples from the Beta distribution. The mean of these samples represents the predicted score, while assessing the variation of the samples provides a measure of uncertainty.

## Conclusions

In this study, we introduced CRISPR-DA, a robust pipeline for CRISPR-Cas9 guide RNA design for detection and diagnosis. Our approach integrates a deep ensemble model, CRISPRDeepEnsemble, to deliver accurate and uncertainty-informed predictions of on-target efficiency.

For off-target analysis, we demonstrated that sequence alignment tools such as BLAST fail to identify all off-target sites and significantly underestimate off-target risk. We confirmed that the Crackling++ off-target scoring module offers superior accuracy and scalability.

Overall, CRISPR-DA addresses several critical challenges in CRISPR-Cas gRNA design by combining cutting-edge bioinformatics approaches with practical, context-specific considerations. It sets a new standard for scalable, accurate, and interpretable gRNA design in the context of detection and diagnosis. CRISPR-DA is available at https://github.com/bmdslab/CRISPR-DA

## Supplementary Note 1: Supplementary Table 1

The full table of viral genomes that were downloaded from the NCBI.

